# Identification of GABAergic neurons innervating the zebrafish lateral habenula

**DOI:** 10.1101/870741

**Authors:** Mahathi Ramaswamy, Ruey-Kuang Cheng, Suresh Jesuthasan

## Abstract

Habenula neurons are constantly active. The level of activity affects mood and behaviour, with increased activity in the lateral habenula reflecting exposure to punishment and a switch to passive coping and depression. Here, we identify GABAergic neurons that could reduce activity in the lateral habenula of larval zebrafish. GAD65/67 immunohistochemistry and imaging of *gad1b:DsRed* transgenic fish suggest the presence of GABAergic terminals in the neuropil and between cell bodies in the lateral habenula. Retrograde tracing with the lipophilic dye DiD suggests that the former derives from the thalamus, while the latter originates from a group of cells in the posterior hypothalamus that are located between the posterior tuberal nucleus and hypothalamic lobes. Two-photon calcium imaging indicates that blue light causes excitation of thalamic GABAergic neurons and terminals in the neuropil, while a subpopulation of lateral habenula neurons show reduced intracellular calcium levels. Whole-cell electrophysiological recording indicates that blue-light reduces membrane potential of lateral habenula neurons. These observations suggest that GABAergic input from the thalamus may mediate inhibition in the zebrafish lateral habenula. Mechanisms governing release of GABA from the neurons in the posterior hypothalamus, which are likely to be in the tuberomammillary nucleus, remain to be defined.

## Introduction

The habenula is a regulator of mood and behaviour in vertebrates (Lecourtier & Kelly, 2007; Namboodiri *et al*., 2016). It consists of two major subdivisions, the medial and lateral habenula, and receives input from several regions of the forebrain (Mok & Mogenson, 1974; Herkenham & Nauta, 1977; Turner *et al*., 2016). The lateral habenula has become a focus of substantial interest recently, as activity here controls the release of broadly acting neuromodulators such as dopamine and serotonin (Wang & Aghajanian, 1977; Hikosaka, 2010; Proulx *et al*., 2014).

In addition to transient activity that is evoked by sensory stimuli (Dreosti *et al*., 2014; Krishnan *et al*., 2014; Cheng *et al*., 2017; Zhang *et al*., 2017) and punishment (Matsumoto & Hikosaka, 2009), the habenula displays persistent or spontaneous activity (Gao *et al*., 1996; Jetti *et al*., 2014; Sakhi *et al*., 2014; Baño-Otálora & Piggins, 2017). This activity is dependent on synaptic input (Kim & Chang, 2005), and changes with time of day (Zhao & Rusak, 2005; Sakhi *et al*., 2014) or as a result of a recent experience. Persistent elevated activity in the lateral habenula is implicated in depression (Li *et al*., 2011; Yang *et al*., 2018; Andalman *et al*., 2019).

Given these effects, there is a growing interest in understanding how activity in the lateral habenula is controlled. In many brain regions, excitation is provided by afferent neurons, with inhibition coming from local GABAergic interneurons. The habenula largely lacks GABAergic neurons (Smith *et al*., 1987; Pandey *et al*., 2018; Zhang *et al*., 2018), but receives GABAergic input (Gottesfeld *et al*., 1979). Although GABAergic input is normally inhibitory, a subset of habenula neurons show extended firing following transient hyperpolarization (Chang & Kim, 2004). In the lateral habenula of mice, afferent neurons from the entopeduncular nucleus and the VTA co-release glutamate and GABA (Root *et al*., 2014; Shabel *et al*., 2014). A small population of GABAergic neurons in the lateral preoptic area (LPO) of rodents projects to the lateral habenula (Barker *et al*., 2017). Another input source to the habenula is the antero-dorsal thalamus (Cheng *et al*., 2017; Zhang *et al*., 2017; Fernandez *et al*., 2018), a region that is rich in GABAergic neurons (Mueller, 2012). In the mouse, GABAergic neurons from the ventral lateral geniculate nucleus and intergeniculate leaflet project to the lateral habenula, and enable light-dependent inhibition (Huang *et al*., 2019). Here we examine the sources of GABAergic input to the lateral habenula of zebrafish, an experimental system with an easily accessible habenula due to eversion of the brain during development (Mueller *et al*., 2011; Folgueira *et al*., 2012). Our data suggests that there are two sources - the thalamus and the posterior hypothalamus. This data on connectivity is expected to shed light on the mechanisms by which brain states are controlled.

## Materials and Methods

### Zebrafish lines

Transgenic lines used include *SqKR11Et (Lee et al., 2010; Teh et al., 2010), Et(−0.6hsp70l:Gal4-VP16)s1020t* (Scott & Baier, 2009), *Tg(UAS:Kaede)s1999t* (Scott *et al*., 2007), *Tg(elavl3:GCaMP6f)a12200* (Wolf *et al*., 2017), *TgBAC(gad1b:DsRed)nns26* (Satou *et al*., 2013), *Tg(vGlut2:GFP)nns43 (Satou et al., 2012), Tg(dao:GAL4VP16)rw0148a* (Amo *et al*., 2014), *Tg(UAS:GCaMP6s)sq205* and *Gt(hspGFFDMC76A)* (Muto *et al*., 2017). The GAL4 enhancer trap line is referred to as *s1020t* in the text for brevity.

All experiments were carried out under guidelines approved by the IACUC of Biopolis (number 191420).

### Neural tracing

A saturated solution of DiD (ThermoFisher Scientific) was made by dissolving a small crystal in 50μL of ethanol. Larval fish were anesthetized with tricaine, mounted in 2% low-melting agarose and placed under a compound microscope (Zeiss Examiner). A small amount of dye was pressure injected into either the left or right habenula by visualizing the fish under a water-immersion 40x objective. The fish was rested for 30 minutes to allow the dye to diffuse and then imaged using a Zeiss LSM800 confocal microscope, under a 40x water-immersion objective. Fish with labelled cells outside the habenula were not used.

### Antibody label

Larvae were fixed in 4% paraformaldehyde in phosphate buffered saline (PBS) overnight and then rinsed several times with PBS. After peeling the skin off the brains, samples were incubated in 1%BSA and then overnight at 4 degrees in primary antibody (anti-GAD65/67, Abcam ab11070, 1:500 (Cheng *et al*., 2017)). The following day the brains were washed several times with PBS and then incubated in secondary antibody (Alexa 488 goat anti-rabbit, 1:1000) overnight at 4°C. After multiple washes with PBS, brains were mounted in low-melting agarose and imaged on a laser scanning confocal microscope (Zeiss LSM800) using a 40x water dipping objective. A total of 7 fish were imaged.

### Electrophysiology

Whole-cell patch clamp and loose-patch recordings were performed from lateral habenula neurons in 5-10 days post-fertilization larvae using procedures previously described in Lupton et al. (2017), except that fish in the current study were not pinned onto a Sylgard dish. Fish were anesthetized by mivacurium before being mounted in a 2% low-melting agarose in a glass-bottom dish that was then immersed with external saline (composition in mM: 134 NaCl, 2.9 KCl, 1.2 MgCl2, 10 Hepes, 10 Glucose, 2.1 CaCl2, pH 7.8). In loose-patch recordings, the pipette was loaded with standard Ringer’s solution to record neural signals from extracellular space while in whole-cell patch recordings, the pipette was loaded with a potassium gluconate–based internal solution (composition in mM: 115 K gluconate, 15 KCl, 2 MgCl2,10 Hepes, 10 EGTA, 4 MgATP, pH 7.2) to record neural signals intracellularly. The pipettes were pulled with thick-walled borosilicate capillaries (1.5 mm OD; 0.86 mm ID; Warner Instruments) using a Flaming Brown P-1000 pipette puller (Sutter Instruments) to give a tip diameter of 1–1.5 μm with an initial resistance of 10–20 MΩ for loose-patch recording. Neural signals were acquired using Multiclamp 700B amplifier, Digidata 1440A digitizer, and pCLAMP software v.10 (all from Molecular Devices). In whole-cell patch recordings, the resistance reached gigaohm seal before breaking into the cells. The data were low-pass filtered at 2 kHz using a Bessel filter and sampled at 20 kHz at a gain of 1. Spike events were detected offline using Clampfit v.10.7 (downloaded from Molecular Devices). Graphs were plotted using Microsoft Excel for whole-cell patch data and in Clampfit for loose-patch data.

### Calcium imaging

Larval fish were imaged as described previously (Cheng *et al*., 2017). Imaging was carried out using a Nikon A1RMP two-photon microscope with a 25x water-dipping objective (NA=1.1), with the Ti-Sapphire tuned to 930 nm and galvano scanning. 5 mm blue LEDs (460-490 nm), controlled by the Nikon Elements software, were used to provide 20 second pulses of light. For imaging the thalamus, multiple planes were captured sequentially.

### Image analysis

Time-series registration, ROI extraction and rastermap calculation were performed using Suite2p (Pachitariu *et al*., 2017). Data was normalized into z-scores using Matlab.

## Results

### The zebrafish habenula is innervated by GABAergic neurons

GAD65/67 immunofluorescence indicates that GABAergic terminals are present in the neuropil of medial (also called dorsal, based on adult morphology (Amo *et al*., 2010)) and lateral (or ventral) habenula of larval zebrafish (Figure 1A, B; arrowheads). In addition, GABAergic neurites were visible between cell soma in the ventro-lateral habenula. The GAD65/67 label in the habenula was distinct from the uniform label seen in cell bodies (see white arrow in Figure 1A), and was seen in the absence of habenula neuron label. As an independent means of visualizing the cell body and neurites of GABAergic neurons, we imaged *gad1b:DsRed, vGlut2:GFP* double transgenic fish, which express DsRed and GFP in GABAergic and glutamatergic neurons respectively. DsRed could be detected in neurites in the lateral habenula, surrounding GFP-expressing neurons (Figure 1C). No DsRed labelled cell was detected in the habenula, in contrast to other brain regions that contained fully labelled cell bodies. This observation, in conjunction with the GAD65/67 antibody labelling, indicates that the zebrafish habenula receives GABAergic input.

**Figure 1.**
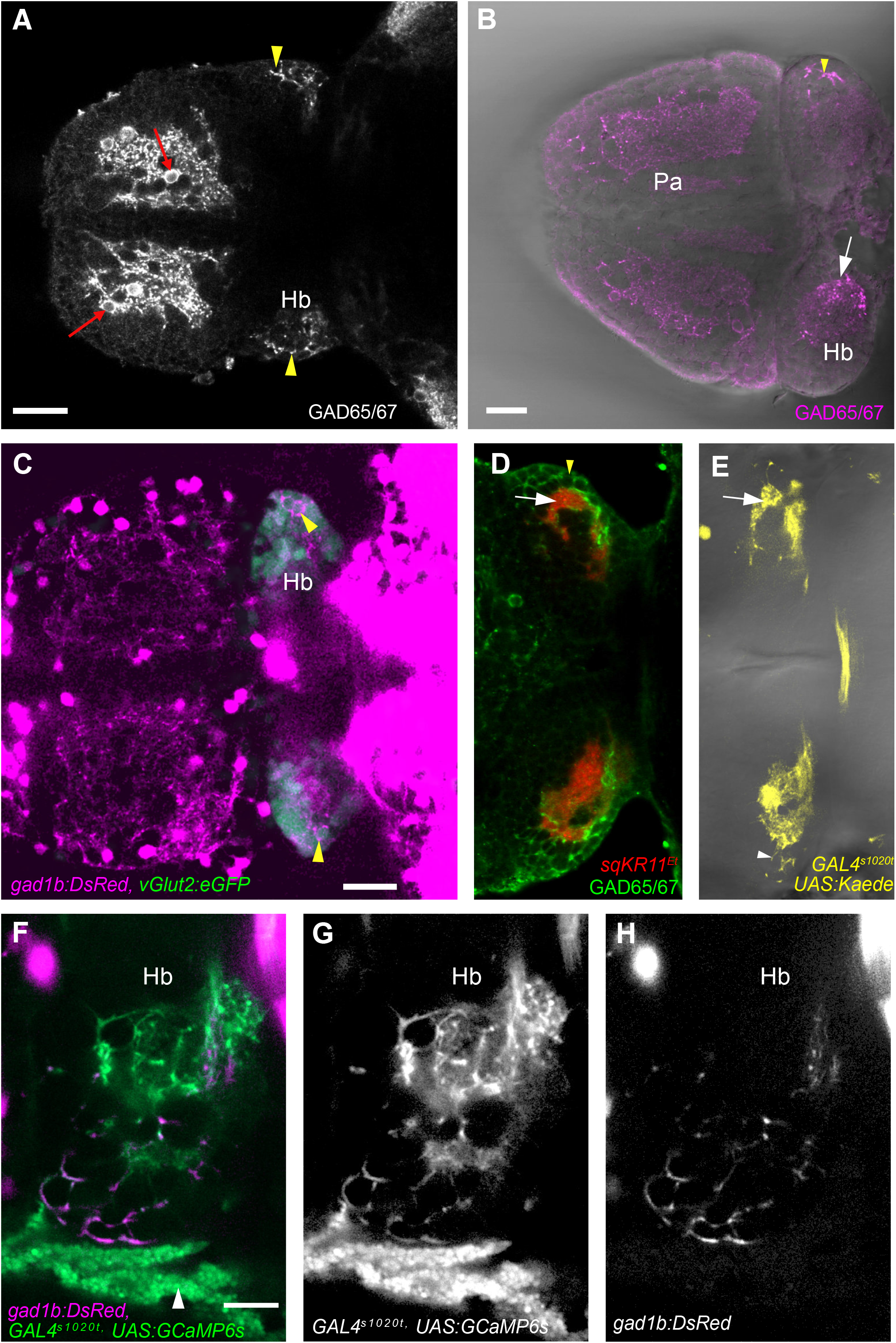
Detection of GABAergic terminals in the habenula. (A) A 6 day old fish, labelled with an antibody to GAD65/67. Terminals are visible in the lateral habenula (yellow arrowheads). The red arrows indicate GABAergic neurons in the pallium. (B) A 2 week old fish, showing GAD65/67-labelled puncta in the neuropil of the medial habenula (white arrow), and in between cells in the lateral habenula (yellow arrowhead). No habenula cell bodies are labelled. This sample is slightly tilted. (C) A 7 day old *Tg(vGlut2:GFP), Tg(gad1b:DsRed)* double transgenic fish. DsRed (magenta) is visible in terminals in the lateral habenula (arrowheads), while habenula cell bodies express GFP (green). (D) An 8 day old fish *SqKR11Et* fish, with afferents from the entopeduncular nucleus labelled in red. Terminals are restricted to neuropil of the lateral habenula (white arrow), and do not overlap with GAD65/67 label between cells (yellow arrowhead). (E) Afferents from thalamic neurons, labelled with Kaede under the control of the *s1020tEt* driver. Terminals are visible in the neuropil (arrow) and also between cells (arrowhead). (F-H) The left habenula of a 6 day old fish, with terminals labelled by GCaMP6s, which is expressed under the control of the *s1020tEt* driver (G), and DsRed under the control of the *gad1b* promoter (H). The white arrowhead in F indicates autofluorescing pterinosomes in xanthophores (Le Guyader & Jesuthasan, 2002). Scale bar = 25 μm for A-E, 10 μm for F-H. All images are dorsal view, with anterior to the left.

### Neuronal tracing identifies sources of GABAergic afferents targeting

To identify the source of the GABAergic inputs, we examined transgenic lines in which habenula afferents are labelled. Two major sources of input to the zebrafish habenula are the entopeduncular nucleus (Turner *et al*., 2016) and the thalamus (Cheng *et al*., 2017). The entopeduncular nucleus is located in the ventral telencephalon, in the vicinity of the lateral forebrain bundle (Turner *et al*., 2016). Neurons in this location, which are labelled in the *Et(sqKR11)* line (Lee *et al*., 2010) (Supplementary Movie 1), were seen to innervate the neuropil of lateral habenula, but no neurites could be detected in between cell soma in lateral habenula (Figure 1D). In contrast, neurites that are labelled by the *s1020tEt* driver (Supplementary Movie 2), were visible between cells of the lateral habenula (Figure 1E-G). These neurites co-expressed DsRed under the control of the *gad1b* promoter (Figure 1H), indicating that they are GABAergic. GABAergic cell bodies were seen in the thalamus in neurons expressing GCaMP6s under the control of the *s1020tEt* driver, but not in the entopeduncular nucleus, which had few 1020tEt positive cells (Supplementary Movie 2). Thus, GABAergic afferents to the lateral habenula may derive from the thalamus.

To confirm the location of the GABAergic afferents, the lipophilic tracer DiD was injected into the lateral habenula of *Tg(gad1b:DsRed)* fish (Fig 2A; n = 17 fish; see Movie 3 for an example). In all cases, terminals between soma were visible in the contralateral lateral habenula, suggesting that there may be bilateral or contralateral origin of these afferents. Retrogradely labelled neurons were visible in the ipsilateral thalamus in only 12 of these fish (Fig 2B, D-G), and out of 42 labelled thalamic neurons, 16 were GABAergic, as judged by *gad1b:DsRed* expression. The fact that 5 fish did not contain any labelled neurons in the thalamus, despite the presence of terminals in the lateral habenula (Figure 2L; see also Movie 20190422) indicates that there is an additional or different source of the soma-targeted terminals. Retrogradely labelled neurons in the entopeduncular nucleus were DsRed-negative (Fig 2C), indicating that this is not the source.

**Figure 2.**
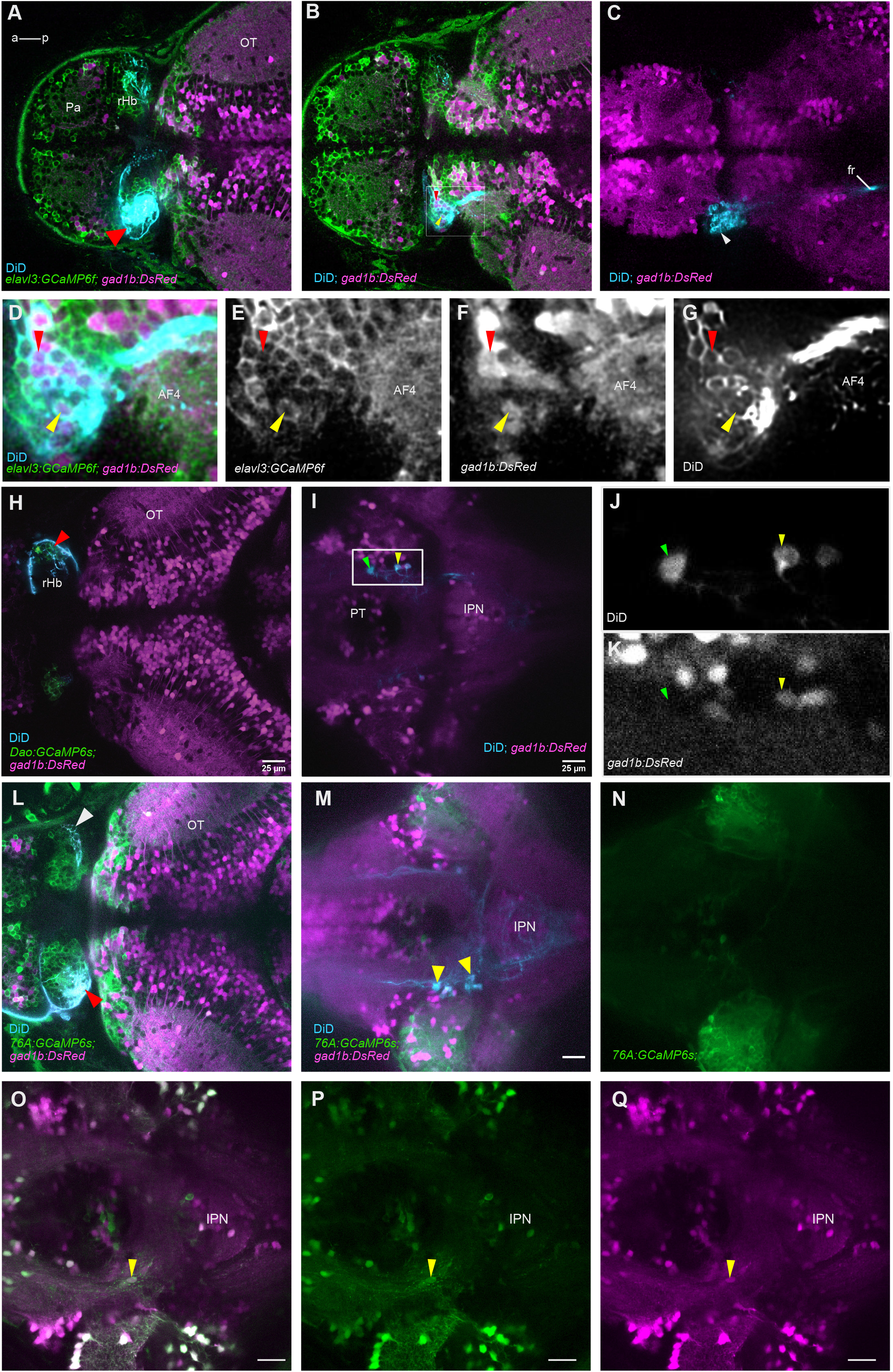
Retrograde label of lateral habenula afferents. (A-C) An example of DiD injection (cyan) into the left lateral habenula (A, red arrowhead) of a 7 day old *Tg(elavl3:GCaMP6f), Tg(gad1b:DsRed)* fish. (B) Retrogradely labelled cell bodies (arrowheads) in the thalamic region. (C) A deeper focal plane. Entopeduncular nucleus (EN) cell bodies are also labelled by DiD but are *gad1b*-negative. (D-G) The boxed area in panel B shown at higher magnification. The arrowheads indicate neurons in the thalamus expressing GCaMP6f (E) and DsRed (F), which have been retrogradely labelled by DiD (H). (H-K) DiD injection (cyan) into the right lateral habenula of another double transgenic 7 day old fish (H, red arrowhead). Lateral habenula neurons express GCaMP6s under the *dao* promoter. (I) Bilateral label of cell bodies in the posterior hypothalamus. (J, K) The boxed region in panel I at higher magnification, showing retrogradely labelled neurons that are *gad1b* positive (yellow arrowhead) or negative (green arrowhead). (L-N) DiD injection into the left lateral habenula of a 7 day old *Tg(76A:GAL4, UAS:GCaMP6s, gad1b:DsRed)* fish. (L) The injection site (red arrowhead). (M) Retrogradely labelled cells (yellow arrowheads), at the level of the inferior lobe of the hypothalamus. (N) GCaMP6s expression in the inferior lobe of the hypothalamus. (O-Q) Expression of GCaMP6s and DsRed in a neuron (yellow arrowhead) in the posterior hypothalamus of 10 day old *s1020t:GAL4, UAS:GCaMP6s, gad1b:DsRed* fish. All panels are dorsal views, with anterior to the left. Pa: pallium, OT: optic tectum, fr: fasciculus retroflexus, rHb: right habenula, PT: posterior tuberculum, IPN: interpeduncular nucleus.

It has been reported that the posterior hypothalamus provides input to the habenula (Turner *et al*., 2016). Indeed, injection into the lateral habenula retrogradely labelled scattered cells in the posterior hypothalamus of all fish where this area was imaged (Figure. 2H-K; n = 7 fish imaged). Label was bilateral in all cases, and 20 out of 46 neurons were positive for DsRed, indicating that this nucleus sends GABAergic projections to the lateral habenula. The labelled neurons were located lateral to the posterior tuberal nucleus, and medial to the inferior lobe of the hypothalamus (IHL), as judged by tracing experiments in the *Tg(76A:GAL4, UAS:GCaMP6s)* line (Muto *et al*., 2017) (Figure 2L-N). Neurons in this region expressed GCaMP6s under the control of the *s1020t* GAL4 driver as well as DsRed under the control of the *gad1b* promoter (Figure 2O-Q). Together, these observations suggest that GABAergic terminals seen between cells in the lateral habenula of the s1020t fish derive from the posterior hypothalamus, and imply that GABAergic terminals in the neuropils derive from the thalamus.

### Blue light triggers activity in thalamic GABAergic neurons and afferents targeting the lateral habenula neuropil

It has been reported previously that cells in the anterior thalamus of zebrafish are reproducibly activated by pulses of blue light (Cheng *et al*., 2017; Zhang *et al*., 2017). To test whether the responding thalamic cells include GABAergic neurons, two photon calcium imaging was performed on larvae expressing GCaMP6s under the control of the s1020t GAL4 driver. An increase in GCaMP6s fluorescence was seen in an average of 46 ± 23 thalamic neurons (mean ± standard deviation; n = 6 fish). Of these, 20 ±13 cells (mean ± standard deviation; approximately 43 ± 18%) were GABAergic (Figure 3A-C). To determine whether terminals in the lateral habenula are also affected, imaging was also carried out in this region (n = 3 fish). In response to blue light, a rise in the fluorescence of this reporter was seen in a subset of terminals within the neuropil of the lateral habenula (Figure 3D-F). No response to blue light was seen in terminals between soma, although activity there was correlated to one another (Figure 3G-I’). These observations are consistent with the notion that blue light can influence the lateral habenula via GABAergic afferents from the thalamus that target the neuropils, but not the cell bodies.

**Figure 3.**
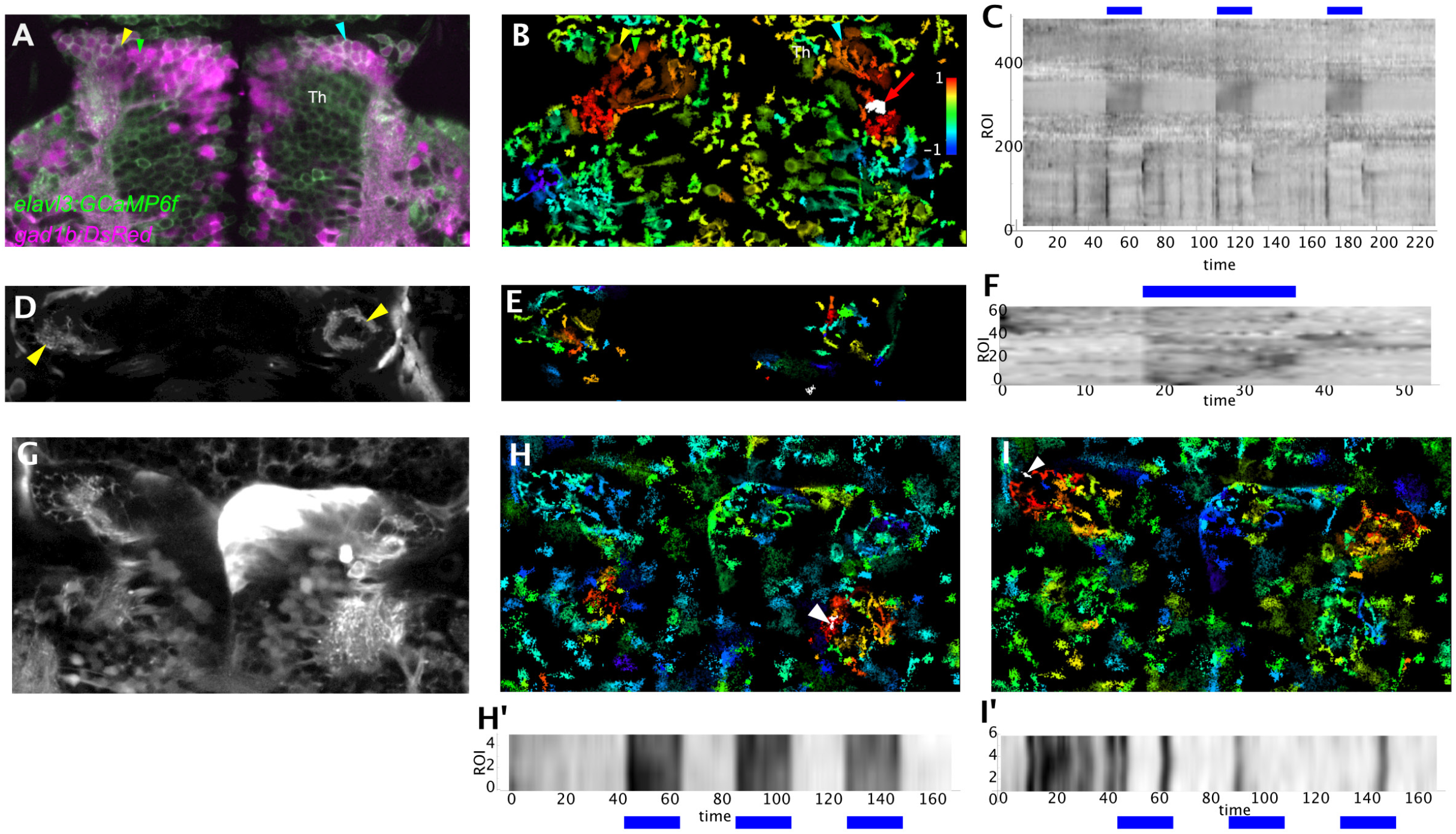
Effect of blue light on thalamic GABAergic neurons. (A-C) The thalamic response of a 6 day old fish to 3 pulses of blue light. (A) Confocal image showing thalamic neurons, some of which express DsRed under the control of the gad1b promoter (magenta). (B) Change in GCaMP6f fluorescence of thalamic neurons, correlated to change in the AF4 neuropil (red arrow), which receives retinal input. Cells indicated by arrowheads have a high correlation with activity in this neuropil and express DsRed, as shown in panel A. (C) Rastermap of cells and neuropils extracted from the time-series. Two major responses are phasic and tonic response to light. (D - F) Calcium imaging of terminals in the neuropil of the ventral habenula of a 5 day old fish exposed to 1 pulse of blue light. (D) Average projection of the time series, showing neuropils (yellow arrowheads). (E) Correlation map of response, with red pixels showing terminals with a highly correlated calcium increase in response to a pulse of blue light. (F) Rastermap showing fluorescence change with time. (G - I) Response in terminals between cell bodies. (G) Average of the time-series, showing the location of terminals (yellow arrowheads). (H) Map of response, correlated to terminals in the thalamic neuropil AF4 (white arrowhead). (H’) Rastermap of 5 ROIs that are highly correlated with blue light pulses. (I) Map of response correlated to terminals in the lateral habenula (white arrowhead). (I’) Rastermap of 6 ROIs in the lateral habenula that are highly correlated to one another. These have no relationship to the light pulses. All panels are dorsal views, with anterior to the top. Th, thalamus. The wedge in panel B indicates the lookup table used for correlation maps.

### Blue light transiently reduces intracellular calcium in a subset of lateral habenula neurons

If thalamic neurons provide GABAergic input to the neuropil of lateral habenula, then stimulation of these neurons may inhibit the lateral habenula. Previous two photon calcium imaging has shown that blue light causes both increase and decrease in ongoing activity throughout the habenula (Cheng *et al*., 2017; Zhang *et al*., 2017). In these recordings, however, it was not clear if inhibition also occurs in the lateral habenula. To determine this, we imaged fish expressing GCaMP6s under the control of the *dao* promoter (n = 6 fish). A minority of cells (30/154) showed a loss of activity during exposure to light (Figure 4A-D). We also performed electrophysiology on intact animals. Both increase and decrease in membrane potential (Figure 4E, F) was detected in lateral habenula neurons by whole-cell recording. When loose-patch recording was employed, the onset of blue light was seen to either trigger or dampen neural spikes (Figure 4G, H). Hence, in addition to excitation, blue light-dependent reduction in the activity of lateral habenula cells is supported by both calcium imaging and electrophysiological recordings.

**Figure 4.**
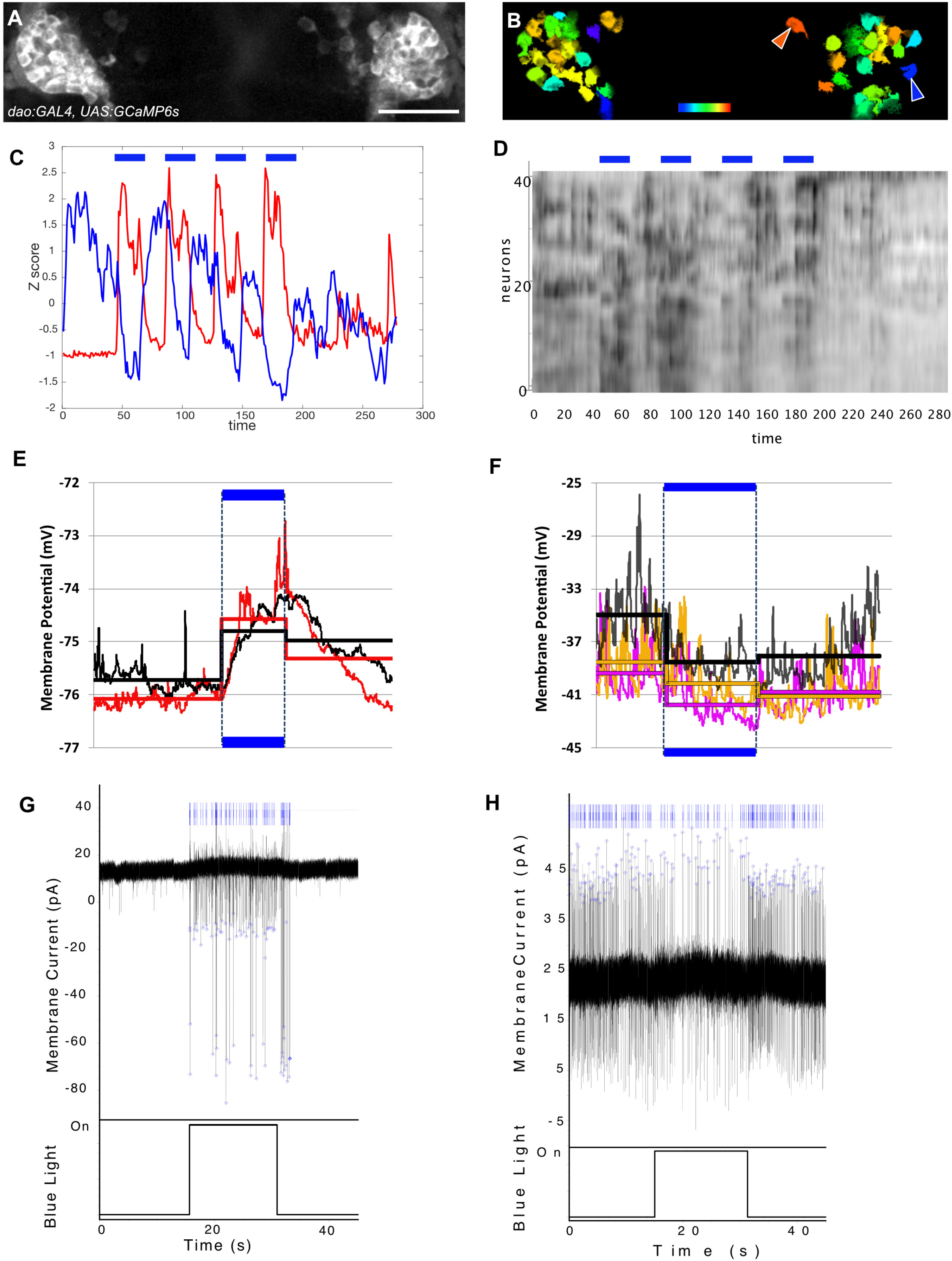
Calcium imaging and electrophysiological recordings in the lateral habenula. (A-D Response of neurons in the lateral habenula of a 5 day old fish exposed to four pulses of blue light. (A) Average of a time-series, showing distribution of cells labelled in a *dao:GAL4, UAS:GCaMP6s* fish. (B) Correlation map of response to 4 pulses of light, with red showing a highly correlated cell, and blue showing an anti-correlated cell. (C) z-scores of the two cells indicated by the blue and red arrowheads in panel B. (D) Rastermap of the response of all cells. Blue bars the presence of blue light. (E, F) Whole-cell recordings showing membrane depolarization (E) or hyperpolarization (F) by blue light. The blue light was 13 seconds in duration in both cases and the range of voltage change (peak-to-trough) was on average 2-3 mV in both cases. Solid horizontal lines represent the means of the membrane potentials before, during and after the blue light in accordance with the trace colors. Each trace in panel E is a separate cell. (G, H) Loose-patch recordings. (G) There are spikes of two different heights, implying signals from two cells. Both show an increase in spikes in the presence of blue light. (H) A recording in another fish, showing a reduction in spiking during blue light delivery. The ticks in the upper trace in (G) and (H) were taken from spike detection from the main trace, as indicated by the blue squares. Scale bar = 25 μm.

## Discussion

We have examined the source of neurons that can influence neural activity in the lateral habenula of larval zebrafish through the release of GABA. GAD65/67 labelling and imaging of a transgenic line which expresses DsRed under the control of the *gad1b* promoter indicate that GABAergic terminals can be found in the habenula neuropils and also in between cell soma. This suggests that GABAergic neurons target both dendrites and the soma of lateral habenula neurons. Targeting onto the soma was not seen in the medial habenula of larval zebrafish, but only in the lateral habenula. Thus, while GABAergic neurons may influence the effect of excitatory input throughout the habenula by inhibition of dendrites, the lateral habenula could experience stronger inhibition due to signalling onto the soma. Tracing data indicates that GABAergic innervation targeting the neuropil and soma derive from different populations.

A recent study in mice demonstrated that GABAergic neurons from the lateral geniculate nucleus innervate and inhibit the lateral habenula in mice (Huang *et al*., 2019). Consistent with these findings, we see that retrograde labelling of lateral habenula neurons in the zebrafish reveals *gad1b*-positive afferents from the thalamus. Further, we show that GABAergic thalamic neurons as well as terminals in the lateral habenula are excited by light, while a number of lateral habenula cells are inhibited. Electrical recordings of the lateral habenula indicate that the onset of blue light led to a reduction in membrane potential and in the number of spikes. The link between illumination levels and mood has been well-studied in animals and 1 humans (Viola *et al*., 2008; Vandewalle *et al*., 2010; Bedrosian *et al*., 2011; LeGates *et al*., 2014). Given that increased activity in the lateral habenula has negative effects (Lecca *et al*., 2014; Proulx *et al*., 2014; Lawson *et al*., 2017; Howe & Kenny, 2018), inhibition of these ^1^ neurons by thalamic GABAergic neurons may contribute to the positive effects of light (Huang *et al*., 2019). However, it should be noted that only a subset of neurons were inhibited by light, indicating that the effect of blue light on inhibition of the lateral habenula, at least in the conditions used here, are relatively modest.

Retrograde tracing experiments also revealed *gad1b*-positive inputs from a cluster of cells lateral to the posterior tuberal nucleus and medial to the inferior lobe of the hypothalamus. Projections from the posterior tuberculum (PT) to the habenula have been previously reported (Hendricks & Jesuthasan, 2007; Turner *et al*., 2016). However, neurons in the posterior tuberculum have been characterized as largely non-GABAergic (Filippi *et al*., 2014; Heap *et al*., 2017). Similar scattered afferents have been identified by retrograde label in the habenula of adult zebrafish (Turner *et al*., 2016), where these were suggested to be located in the posterior hypothalamus, and in the trout (Yañez & Anadón, 1996), where the cells were named the “tuberculohabenular nucleus”. We propose that these cells are located in the tuberomammillary nucleus, which is located in the posterior hypothalamus (Chen *et al*., 2017). This would be consistent with the existence of histaminergic fibers in the lateral habenula (Kaslin & Panula, 2001), the fact that neurons in the tuberomammillary nucleus in the posterior hypothalamus are the sole source histamine in vertebrates, and that histaminergic neurons co-release GABA (Sundvik & Panula, 2012; Yu *et al*., 2015). Future experiments to define the function of these cells should yield fresh insight into the mechanisms by which activity in the lateral habenula is inhibited.

## Acknowledgements

This research was supported by a Lee Kong Chian School of Medicine, Nanyang Technological University Singapore Start-Up Grant, and the Singapore Ministry of Education under its Academic Research Fund Tier 2 (MOE2017-T2-058) and Tier 1 (MOE2016-T1-001-152) awards. We thank Vatsala Thirumalai and members of her lab for instruction on techniques in zebrafish electrophysiology.

## Author contributions

MR performed antibody label, retrograde tracing and microscopy. RKC performed electrophysiology and functional imaging. SJ imaged transgenic lines, analyzed data and wrote the manuscript.

## Data Accessibility

All confocal stacks used here are available on Figshare.

## Supplementary Material

**Supplementary Movie 1. Z-stack of a 5 day old SqKR11Et, elavl3:GCaMP6f fish.** The yellow arrowheads indicate the lateral forebrain bundle in the ventral telencephalon. The white arrowhead indicates cells in the entopeduncular nucleus, which project to neuropils in the habenula.

**Supplementary Movie 2. Z-stack of a 6 day old Tg(s1020tGAL4, UAS:GCaMP6s, gad1b:DsRed) fish.** The yellow arrowheads indicate the lateral forebrain bundle in the ventral telencephalon. The white arrowhead indicates the entopeduncular nucleus, which contains no DsRed expressing cells. Cells in the thalamus are strongly labelled by the s1020t driver.

**Supplementary Movie 3. Retrograde label of cells in the posterior hypothalamus.** Z-stack of a Tg(dao:GAL4, UAS:GCaMP6s, gad1b:DsRed) fish, after DiD label of the left lateral habenula. Labelled cells are visible in the posterior hypothalamus. Habenula projections to the raphe are also visible. There are no projections to the IPN. Note that GCaMP6s is expressed in the posterior hypothalamus.

